# Targeted Drug Delivery by Radiation-Induced Tumor Vascular Modulation

**DOI:** 10.1101/268714

**Authors:** Sijumon Kunjachan, Shady Kotb, Rajiv Kumar, Robert Pola, Michal Pechar, Felix Gremse, Reza Taleeli, Florian Trichard, Vincent Motto-Ros, Lucie Sancey, Alexandre Detappe, Andrea Protti, Ilanchezhian Shanmugam, Thomas Ireland, Tomas Etrych, Srinivas Sridhar, Olivier Tillement, G. Mike Makrigiorgos, Ross Berbeco

## Abstract

Effective drug delivery is severely restricted by the presence of complex pathophysiological barriers in solid tumors. In human pancreatic adenocarcinoma, mature and hypopermeable tumor blood vessels limit the permeation and penetration of chemo or nanotherapeutics to cancer cells and substantially reduce the treatment efficacy. New, clinically-viable strategies are therefore sought to breach the neoplastic barriers that prevent optimal tumor-specific drug delivery. Here, we present an original idea to boost targeted drug delivery by selectively knocking down the tumor vascular barrier in a poorly permeable human pancreatic cancer model. For the first time, we demonstrate that clinical irradiation (10 Gy, 6 MV) can induce tumor vascular modulation when combined with tumor endothelial-targeting gold nanoparticles. *Active* disruption of tumor blood vessels by nanoparticle-combined radiotherapy led to increased vessel permeability and improved tumor uptake of two prototypical model nanodrugs: i) a short-circulating nanocarrier with MR-sensitive gadolinium (Gad-NC; 8 kDa; t_1/2_=1.5 h) and ii) a long-circulating nanocarrier with fluorescence-sensitive NIR dye (FL-NC; 30 kDa; t_1/2_=25 h). Functional changes in the altered tumor vessel dynamics, measured by relative changes in permeability (*K*_*trans*_), flux rate (*K*_*ep*_) and extracellular interstitial volume (*V*_*e*_) were consistent with the concomitant increase in nanodrug delivery. This combination of radiation-induced antivascular and nanodrug-mediated *anti-tumor* treatment offers high therapeutic benefit for tumors with pathophysiology that restricts efficient drug delivery.

GRAPHICAL ABSTRACT

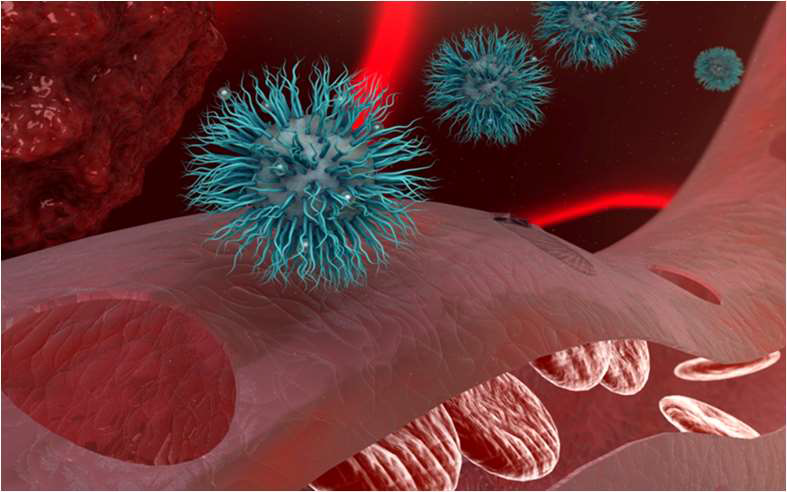

---

Tumor-specific drug delivery is limited by the presence of several pathophysiological barriers that often lead to sub-optimal tumor accumulation. ^1–6^ Clinical experience with nanodrugs, including routinely used Doxil (Johnson & Johnson), Abraxane or *nab-Paclitaxel* (Celgene), demonstrated substantial progress in the minimization of drug toxicity, albeit with little therapeutic gain. While mounting experimental evidence continues to address this issue, recent analyses show that the overall accumulation of tumor-targeted nanomedicines in solid tumors is mere ~1-2%. ^5,7^ Moving forward, the far-reaching potential of nanomedicines will remain unrealized unless there is a progressive effort in identifying and overcoming the physiological barriers that prevent optimal cancer drug delivery. ^5, 8–10^

Tumor neovasculature is a key barrier to drug delivery and a prime target for chemo and radiation therapy (RT). ^11–18^ There are reports of mixed clinical success with chemical vascular disrupting agents (cVD) such as combretastatin or ombrabulin, which cause pulmonary embolism, coronary vasospasm, and other cardiovascular toxicities. ^15, 19^ We have devised strategies where (nontoxic and) tumor blood vessel-targeted gold nanoparticles in combination with RT can be used as vascular disrupting agents. ^16, 20–21^ Thereby, the inadvertent toxicities associated with the use of cVDs can be considerably reduced, leading to better cancer therapy.

High-Z metallic nanoparticles provide a local radiation boost during RT due to the increased photoelectric interactions. ^20–25^ In optimal doses, gold nanoparticles are safe, biocompatible and clinically useful. ^5–6 26–33^ Ultrasmall nanoparticles of 1-5 nm are renally eliminated, and long-term clearance studies show that gold nanoparticles are eliminated from the body *via* phase degradation mechanisms without invoking chronic adverse reactions. ^34–35^ Vascular targeting ligands such as cRGD (a cyclo-pentapeptide) have a strong binding affinity to the α_v_β_3_ and α_v_β_5_ integrin receptors present along the tumor endothelial linings. ^16^ When attached to PEGylated-gold, prominent localization along the pancreatic tumor blood vessels has been observed. Moreover, the dense stromal matrix of human pancreatic adenocarcinoma tumor model (*h*-PDAC) leads to the perivascular retention of nanoparticles, contributing to indirect radiation responses. ^16^

Tumor endothelial targeted and long-circulating gold nanoparticles (t-NP) can cause tumor vascular disruption under set irradiation conditions in pancreatic tumors. ^16^ The sub-mm precision of modern clinical radiation therapy (RT) and the target-specificity of t-NP render it a “dual-targeted” treatment with high spatiotemporal accuracy. We hypothesized that the local induced damage to the tumor neovasculature might lead to improved vascular permeability and enhance tumor-specific drug delivery. This is particularly significant in *h*-PDAC due to its poor vascular permeability and low uptake of anticancer drugs or nanotherapeutics. ^5^

We demonstrate that targeted drug delivery can be deployed by the selective radiation amplification induced damage to the tumor neovessels in *h*-PDAC. Noninvasive MRI and fluorescence studies using short and long-circulating nanocarriers demonstrate an increase in: i) quantitative tumor uptake, ii) tumor vascular permeability, and iii) intratumoral distribution of polymeric nanomedicines in *h*-PDAC.

## RESULTS AND DISCUSSION

Passive tumor targeting relies on the inherent defects of tumor blood vessels. ^36^ The rapid and angiogenic tumor growth tumor leads to hyperpermeable and defective endothelium that permits the transport of nanocarriers across the blood vessels. Furthermore, the absence of a fully functional lymphatic system further aids in the retention of nanoparticles within the tumors. This phenomenon, known as the “enhanced permeability and retention” (EPR) effect, forms the basis for the accumulation of most clinically approved nanomedicine formulations. ^37–38^ EPR is highly variable across various tumor models. ^6, 39^ Our previous studies show that the inter and intratumoral heterogeneity of EPR is directly related to the angiogenic profile and growth rate of the tumors. ^6, 16, 38^ In contrast, slow-growing tumors (ex. *h*-PDAC) show intact, mature vessels (adequately sheathed by α-SMA or pericytes) that has a high receptor expression density on its tumor neoendothelium. Hence, the passive accumulation of anticancer or nanodrugs is relatively low in h-PDAC tumors. ^16, 38, 40–41^ To breach the tumor vascular barrier in *h*-PDAC, we have utilized gold nanoparticles conjugated with RGD to anchor to the tumor neoendothelium, and by using external beam-RT, tumor vascular disruption was induced. The transient alteration in tumor blood vessel facilitated enhanced and tumor-specific drug delivery (**Fig. 1**).

**Figure 1:**
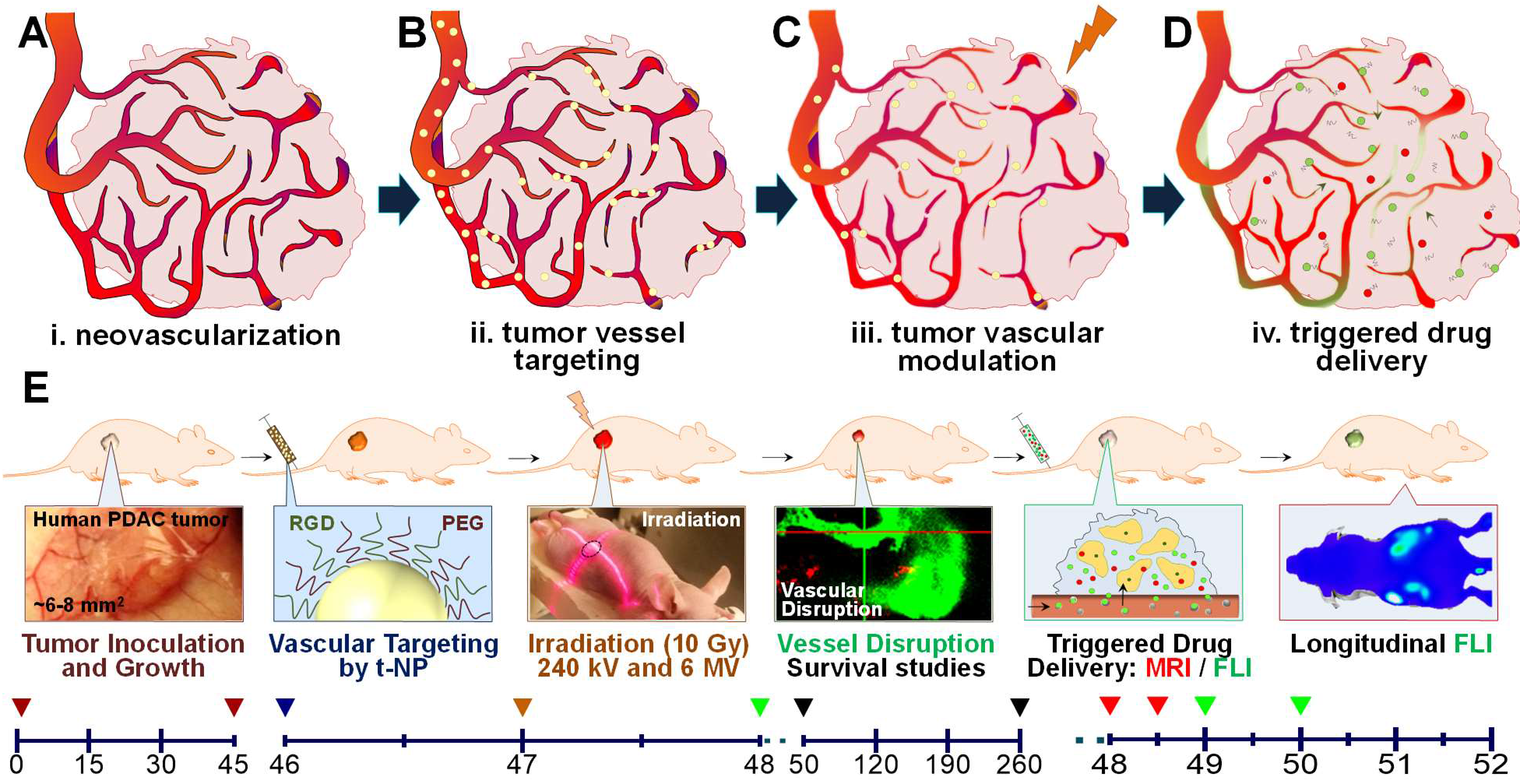
Concept and study design. A schematic depiction of radiation-induced tumor vascular modulation to trigger a tumor-specific drug delivery. **A-D**. In an angiogenic human pancreatic adenocarcinoma tumor, gold nanoparticles (in yellow) were targeted to the α_v_β_3_, α_v_β_5_ integrin receptors present along the tumor neovessels, and specific radiation damage is induced. This results in selective vascular rupture and leads to the triggered delivery of tumor-specific polymeric nanocarriers. Short-circulating MR-sensitive (red; Gad-NC) and long-circulating fluorescence-sensitive (green; FL-NC) polymeric nanocarriers were tested for improved drug payload delivery. **E**. Experimental procedures and timelines are shown in days. Following tumor inoculation and growth (day 0 - 45), the study was carried out in three stages. Phase I: Inducing selective radiation damage to pancreatic tumor neoendothelium (day 46 - 48); Phase II: Imaging tumor vascular disruption and assessing the survival benefits (day 50 - 260); Phase III: Triggering tumor-specific payload delivery of Gad-NC and FL-NC - two representative model nanodrugs (day 48 - 52). Furthermore, this study involved the use of targeted gold nanoparticles (t-NP); preclinical and clinical irradiations (240 KV and 6 MV); T1-weighted and DCE-MR and whole-body fluorescence imaging using Gad-NC and FL-NC respectively at stipulated time-points. For all the survival, treatment or imaging studies, a tumor size of ≥ 2 cm was considered as a terminal endpoint.

### Physicochemical characterization and *in vitro* testing of targeted nanoplatform

Heterobifunctional, PEG/RGD-modified gold nanoparticles (t-NP) were synthesized based on the standard turkevich method. ^16, 20, 42^ With spherical morphology, monodisperse t-NP show a core size of 2-3 nm and hydrodynamic size of 5-10 nm (**Fig. 2A-B**). Tumor endothelial targeting was accomplished *via* RGD - a standard vascular targeting ligand that docks to the transmembrane receptor proteins (α_v_β_3_ and α_v_β_5_) present along the vascular lumen ^16, 38, 43^. Preliminary simulation studies show linear differences in the electron emission spectra with varying particle sizes of t-NP. An optimal core size of ~2-3 nm predicted the highest fluence of emitted electrons and the subsequent photoelectric interactions (**Fig. 2C**). Further analysis of DNA double-strand breaks (D-DSB) using Monte Carlo Damage Simulation (MCDS) studies confirmed an increase in D-DSB due to the specific nanoparticle-radiation interactions (supplementary, **Table 1**).

**Figure 2:**
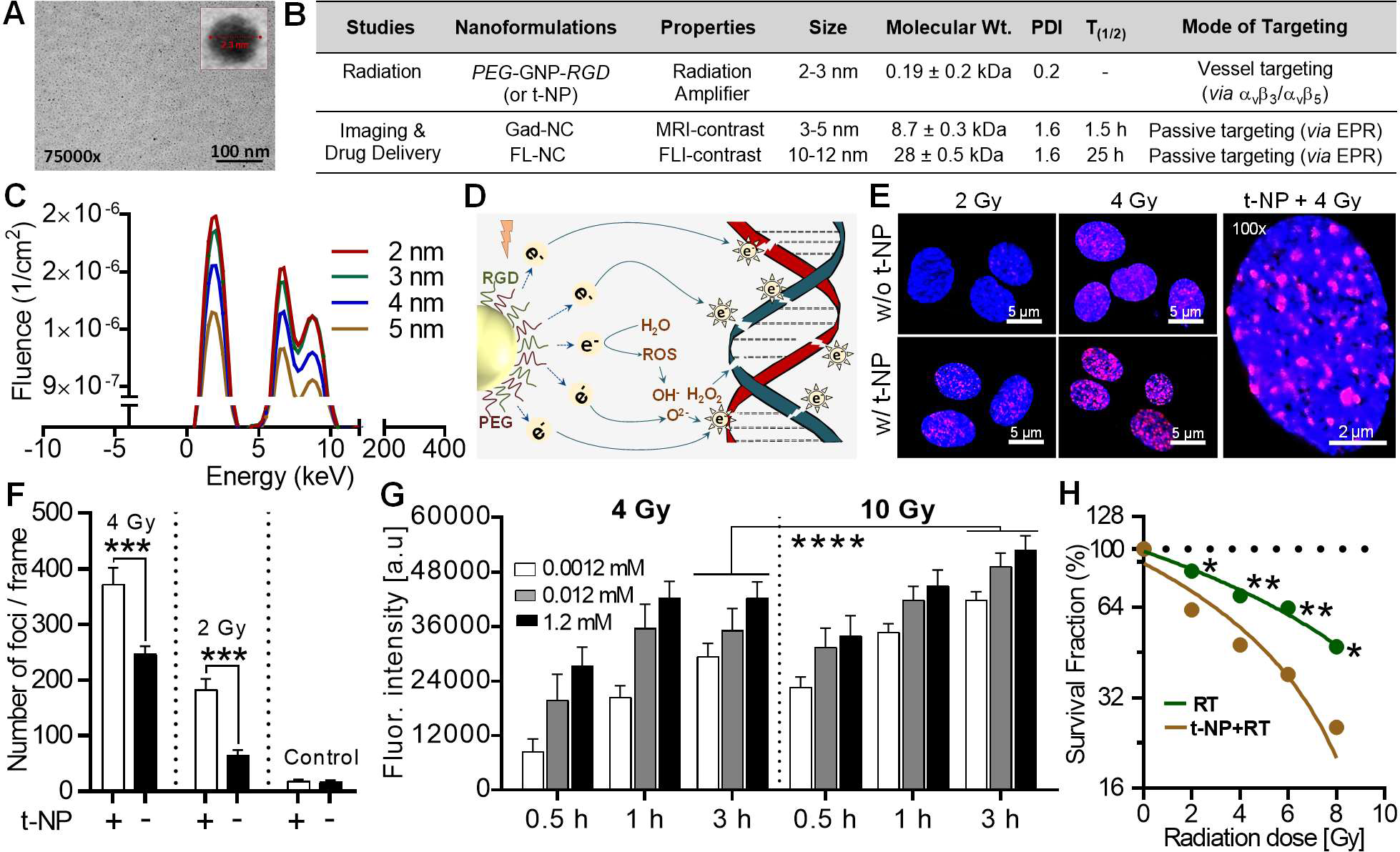
Physicochemical characterization and *in vitro* radiation damage amplification. **A**. High-resolution TEM image shows ultrasmall gold nanoparticles with a core size of 2-3 nm (cf. inset) bi-functionalized with *Arg-Gly-Asp* (RGD) and PEG (polyethylene glycol). **B**. The table summarizes various nanoformulations used in this study: targeted gold nanoparticles (t-NP) mediated a radiation-specific tumor vascular disruption; MR and fluorescence-contrast polymeric nanocarriers with diverse physicochemical properties were used for EPR-mediated triggered drug delivery studies. **C**. Preliminary simulation studies showed the relationship between the ejection of low energy electrons from the gold nanoparticles and the respective size. Gold nanoparticles (t-NP), with a core size of 2-3 nm is predicted to generate superior radiation dose amplification due to the reduced self-absorption of Auger electrons. **D**. Schematic illustration of physical and biological radiation interactions that lead to DNA double-strand breaks (D-DSB). Low energy electrons generated due to the radiosensitization of t-NP induce direct D-DSB, and the simultaneous generation of free radical’s results in an indirect DNA damage. **E-F**. DNA damage studies following radiation and (+/−) t-NP treatment shows distinct differences (~twofold) in the D-DSB in proliferating human endothelial cells. Further quantification of the damaged foci confirmed significant differences between nanoparticle-treated and non-treated groups under different irradiation conditions. **G**. Free radical assays (primarily for peroxides) at three different t-NP concentrations (0.0012, 0.12, and 1.2 mM) showed dose-dependent changes in the free-radical mediated damage to HUVEC at different time point’s post-RT. The change in fluorescence signal intensity corresponds to the amount of reactive oxygen species detected. The data were normalized to the non-treated control: 0 Gy and without t-NP. **H**. Linear, quadratic regression plot of endothelial cell survival demonstrated significant differences at 2 Gy (*P*=0.018), 4 Gy (*P*=0.009), 6 Gy (*P*=0.006) and 8 Gy (*P*=0.011) in the t-NP+RT vs. the RT-only treatment. All results were normalized to the respective treated and non-treated controls. Of note, error bars are smaller than the dotted plots.

Radiation dose amplification is mediated by direct physical damage due to the emission of low-energy electrons that cause D-DSBs and indirect biological damage due to the release of OH^−^ (hydroxyl), H_2_O_2_ (peroxide) and O^2−^ (superoxide anions) based ionizing radicals (**Fig. 2D**). ^44^ The radiation response of proliferating human umbilical endothelial cells at 2 and 4 Gy demonstrated substantial cellular damage in combination with t-NP. More than two-fold increase in D-DSB was measured (**Fig. 2E-F**). Free radical generation from 15 min to 3 h post-RT at various t-NP concentrations showed that both 4 and 10 Gy improved the free radical (primarily peroxide) mediated radiation damage in the t-NP+RT group compared to its ‘no nanoparticle’ treatment controls (**Fig. 2G**). Higher concentrations of gold nanoparticles can affect the cell proliferation by inducing changes in cell morphology and toxicity effects, and therefore we optimized a sub-toxic dose that is suitable for *in vitro* purposes (**Fig. S1**).^45^ Direct clonogenic response studies illustrate that t-NP caused radiosensitization in proliferating endothelial cells with a sensitivity enhancement ratio (SER) of 1.35. Overall, t-NP+RT demonstrated significant cell damage at different radiation doses: 2 Gy (*P*=0.017), 4 Gy (*P*=0.008), 6 Gy (*P*=0.06) and 8 Gy (*P*=0.0112), compared to the RT-only group (**Fig. 2H**). *In vitro* studies prove that t-NP in combination with RT results in high endothelial cell damage *via* both physical and chemical mechanisms.

### Biodistribution and tumor localization studies

Following *i.v.* administration of t-NP in *h*-PDAC tumor-bearing mice, progressive and longitudinal intratumoral accumulation was recorded using ICP-MS. Maximum tumor accumulation of 2 - 2.5% ID was measured at 24 h (**Fig.3A**). Further quantification of t-NP in other vital organs showed a considerable decline at around the same time frame. Particularly in the heart and kidneys, t-NP concentrations declined to < 2% ID; non-specific uptake in the liver showed ~ 6% ID at 24 h. The literature reports that gold nanoparticles of size ≤ 10 nm that accumulate in the liver are cleared out of the body *via* phase degradation mechanisms following the hepatobiliary pathways. ^30, 46^

t-NP accumulation in the tumor blood vessels was further confirmed by LIBS (Laser-induced breakdown spectroscopy imaging) - a technique that accurately detects/images endogenous metals and correlates with hyperspectral data in real-time. LIBS imaging captured gold (*Au*) signals from t-NP in the *in vivo* tumor specimens collected at 1 and 24 h post-*i.v*. administration (**Fig. 3B**). A cross-examination of various tumor slices shows the heterogeneous distribution of t-NP from the tumor periphery to the core. In agreement with the biodistribution data, maximum t-NP accumulation was observed at 24 h. Importantly, a strong correlation was noticed between the Au from t-NP and *Fe* from the stagnated blood vessels (**Fig. 3C**). *Au* - *Fe* correlation studies (coefficient ratio=0.65) confirmed their co-localization within the tumor, and particularly along the tumor blood vessels (**Fig. S2**). A spectral peak at 268 nm confirmed the presence of *Au* in the respective (Capan-1) *h*-PDAC tumors (**Fig. 3D**). Furthermore, histological staining of tumor samples (collected at 24 h post-*i.v*. administration of t-NP) and microscopic imaging affirmed the presence of *Au* predominantly along the endothelial walls in *h*-PDAC (**Fig. S3**).

**Figure 3:**
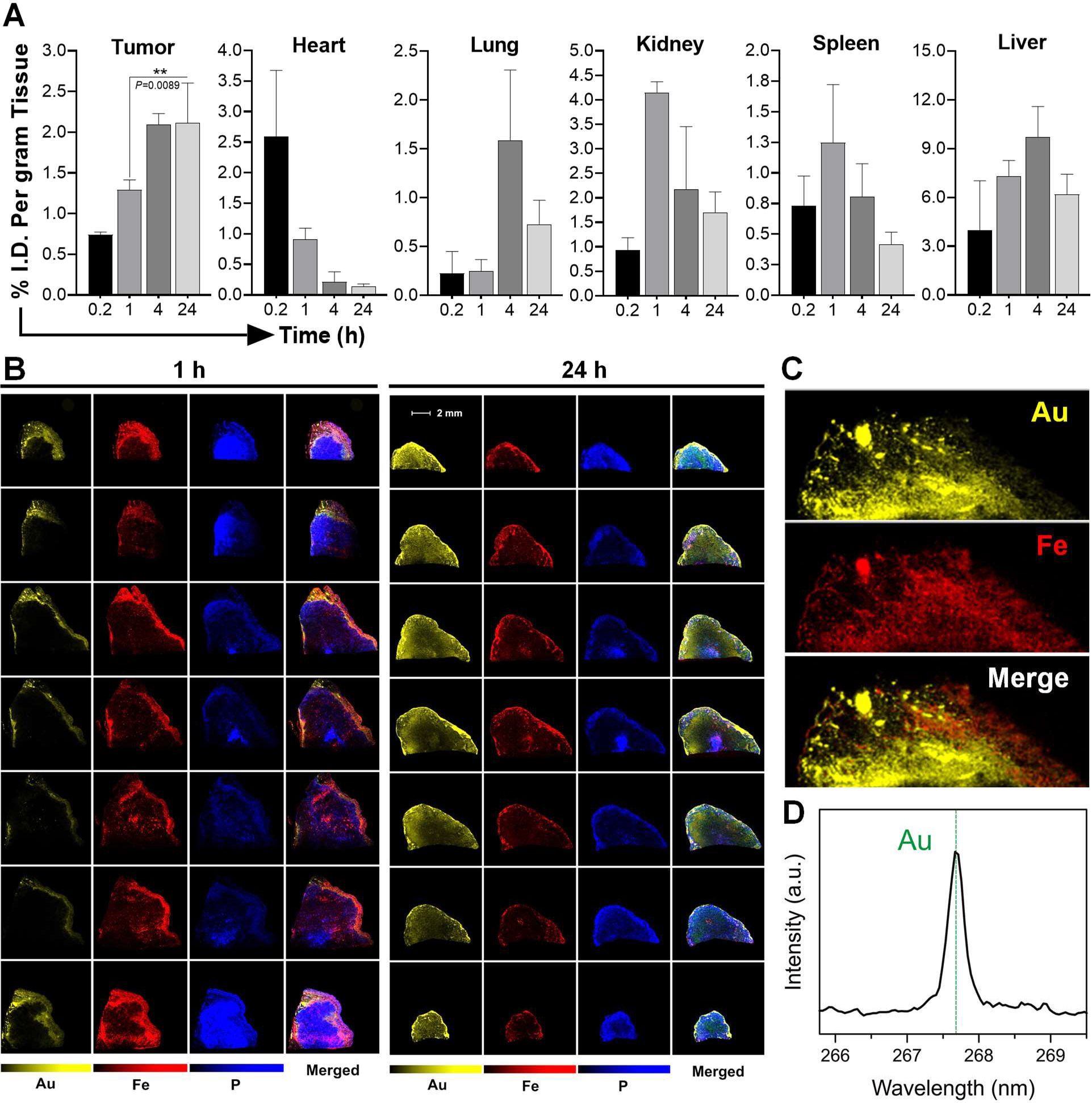
Biodistribution and tumor localization studies. **A.** Quantitative biodistribution of t-NP in tumor and various organs were measured by ICP-MS following its *i.v*.-administration in Capan-1 *h*-PDAC tumor-bearing mice (*n*=5). At 24 h, maximum tumor accumulation was noticed, whereas the distribution of t-NP declined in other organs. **B**. Laser-induced breakdown spectroscopy or LIBS imaging was performed to qualitatively estimate the intratumoral distribution of t-NP. The symbols correspond to *Au* - Gold (indicative of t-NP); *Fe* - Iron (a surrogate marker for tumor blood vessels); *P* - Phosphorus. The presence of *Au* from t-NP and *Fe* from the heme of blood cells gave specific signals in LIBS imaging. A complete tumor analysis from the periphery to the core show heterogeneous distribution of nanoparticles. 24 h tumor samples show the maximum nanoparticle (or *Au*) accumulation in the tumor specimens, and often seen in proximity to the tumor blood vessels (red). **C**. High-magnification LIBS image shows substantial overlap of *Au* with *Fe* in a 24 h tumor sample. **D**. The real-time spectral analysis demonstrated a corresponding peak for *Au* (yellow) at 267.595 nm and confirmed its intratumoral accumulation in an *h*-PDAC tumor model.

### Tumor vascular modulation and survival studies

Both biodistribution and microscopy data confirmed 24 h as an ideal time-point for RT due to the maximum tumor: clearance ratio. *In vivo* studies show that preclinical-RT combined with t-NP demonstrated significant anti-vascular effects (**Fig. S4**). In a *h*-PDAC tumor model (Capan-1), extended survival of ≥75 days was observed in the case of t-NP+RT treated (*P*<0.002), compared to RT-only treatment (**Fig. 4A**). Clinical radiation was applied at specific orthogonal angles to the tumors, and optimal tumor coverage with minimal radiation exposure to peripheral organs was attained. Radiation dose distribution data confirmed that ≥99% of the tumor region received an equivalent dose of 10 Gy (6 MV), while the normal tissues were essentially spared (≤1%) (**Fig. 4B**). A survival rate of ~80% was observed in ‘nanoparticle-combined clinical-RT group.’ Interestingly, all of the surviving mice from the ‘t-NP+RT’ group exhibited a complete tumor remission with no signs of toxicity or health issues were observed (**Fig. 4C**).

**Figure 4:**
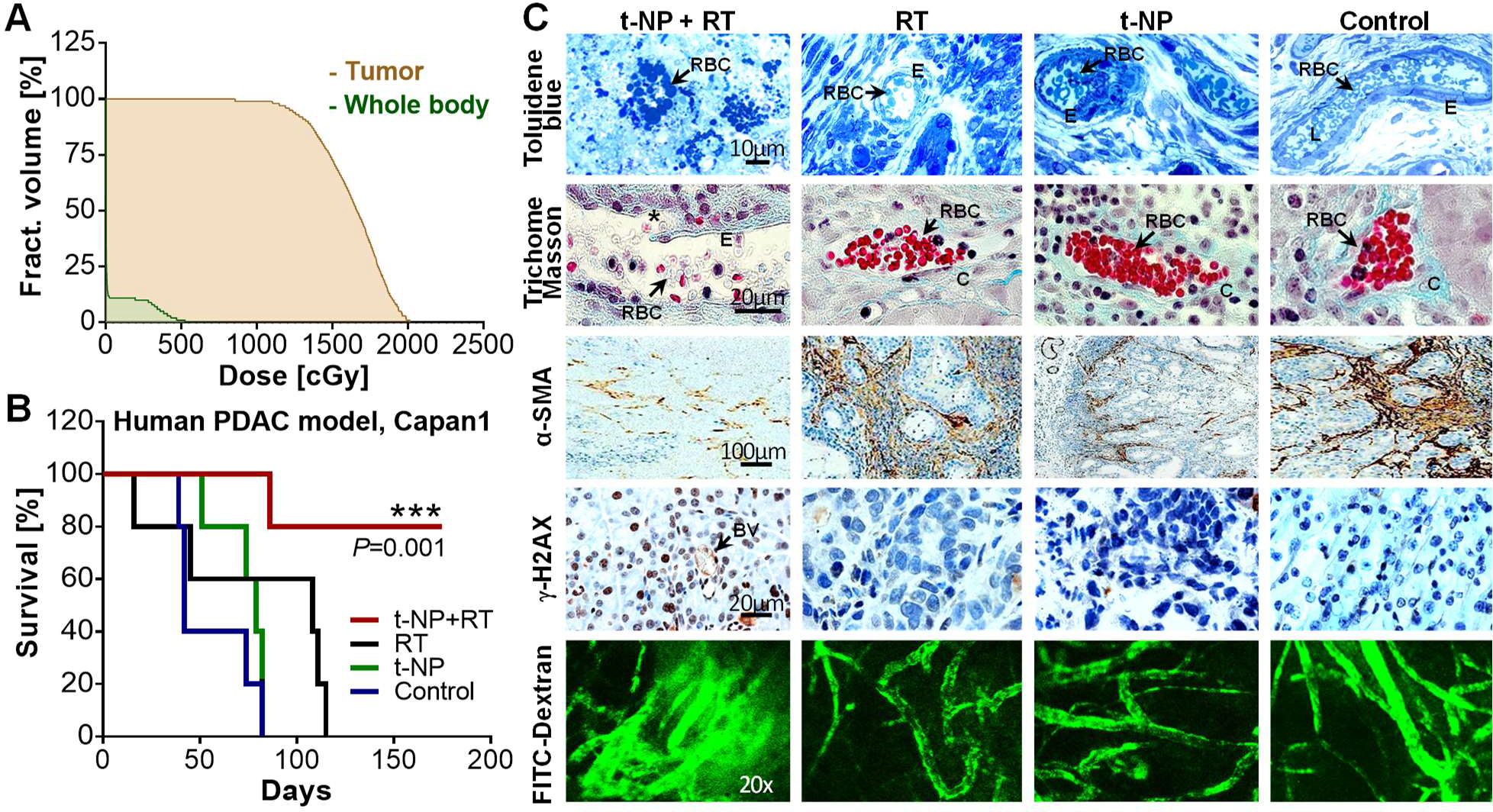
Selective radiation damage and survival studies. Tumor-selective radiation damage elicited changes in the survival and vessel morphology in *h*-PDAC. t-NP was administered intravenously at a dose of 1.2 mg/g, and respective irradiations were carried out at 24 h post-administration. **A**. A dose-volume histogram was recorded to estimate the precise tumor dose delivery compared to the rest of the body in a clinical beam-RT (6 MV, 10 Gy) set-up. Dose distribution calculations indicate that >99% of the tumor region received a radiation dose of 10 Gy. **B**. Kaplan-Meier plot depicting survival studies using clinical beam irradiations (6 MV, 10 Gy) demonstrate an improved therapeutic benefit with the t-NP+RT treatment. Log-rank (Mantel-Cox) tests were used for statistical analysis. **C**. Histological evidence confirmed the vascular damage at 24 h post-treatment using different qualitative techniques: Toluidine blue and trichome Masson staining has confirmed the rupture of tumor neovessels during combined radiation and nanoparticle treatment. Selective rupture (denoted by asterisks in the figure), resulting in non-functional and apoptotic RBC’s (arrows) and vascular instability was observed. However, functionally-viable and collagen-sheathed active vessels were present in other control samples. DNA damage studies using γh2ax confirmed radiation-specific damage in the tumor cells and vessels. Brown color indicates DNA damage during combined t-NP and RT treatment. Smooth muscle actins (α-SMA; brown) that support the tumor endothelium have collapsed during radiation and nanoparticle-induced tumor vascular modulation. Functional (or perfusion) damage at 24 h post-radiation, assessed by the FITC-dextran infusion (70 kDa) studies show extensive leakage (or permeation of FITC) in selective, treated vessels in the t-NP *plus* RT groups. Under all other treatment conditions, the vessels were distinctly labeled, and no signs of passive leakage into the interstitial tumor spaces was evident. RBC: red blood cells; E: endothelium; L: lumen; C: collagen; BV: blood vessel.

The spatiotemporal localization of t-NP and its subsequent exposure to RT induce ‘tumor vascular modulation’ in PDAC tumors. Further histological examinations confirmed that the experimental group receiving t-NP+RT demonstrated a high degree of tumor vascular disruption after clinical-RT (6 MV, 10 Gy). Morphological changes in the tumor neovasculature were evident at 24 h in the ‘nanoparticle combined RT group.’ Loss of endothelial integrity and specific blood vessel rupture were evident in the t-NP+RT group whereas the other controls showed intact and fully functional blood vessels without any apparent damage. Blood cells, primarily RBCs, underwent apoptotic changes in the ‘nanoparticle combined RT group’ when analyzed by trichome or toluidine blue stainings (**Fig. 4D**). The histological evidence further showed that the supporting smooth muscle actins (α-SMA) were largely compromised (**Fig. 4D**). Radiation-specific (γH2AX staining) confirmed massive DNA double-strand breaks (indicated by brown nuclei) under t-NP+RT treatment (**Fig. 4D**). FITC-dextran perfusion based (functional) studies show a considerable loss of vessel integrity, leading to the FITC diffusion through altered tumor blood vessels (**Fig. 4D**). On the other hand, the control groups show intact, non-permeable and functional tumor blood vessels. Together, both the survival and histological evidence confirmed the potential of t-NP as tumor vascular disrupting agents under both preclinical and clinical radiation therapy conditions.

### Enhancing tumor-specific drug delivery

By modulating the tumor vascular barrier, we anticipated an enhancement in tumor-specific drug delivery in *h*-PDAC. To achieve this, we employed two prototypical nanodrug carriers, each with a unique size, circulation, and imaging capabilities: I) A short-circulating MR-sensitive gadolinium nanocarrier (Gad-NC) and II) A long-circulating fluorescence-sensitive HPMA nanocarrier (FL-NC) was used for further studies (**Fig. 5A**). Gad-NC demonstrates rapid systemic circulation (t_1/2_=1.5 h) and is cleared *via* the kidneys relatively faster. ^47–49^ With a molecular weight of ~8.7 kDa, Gad-NC has a maximum tumor uptake at 15-30 min post-*i.v*. administration. ^50–51^

**Figure 5:**
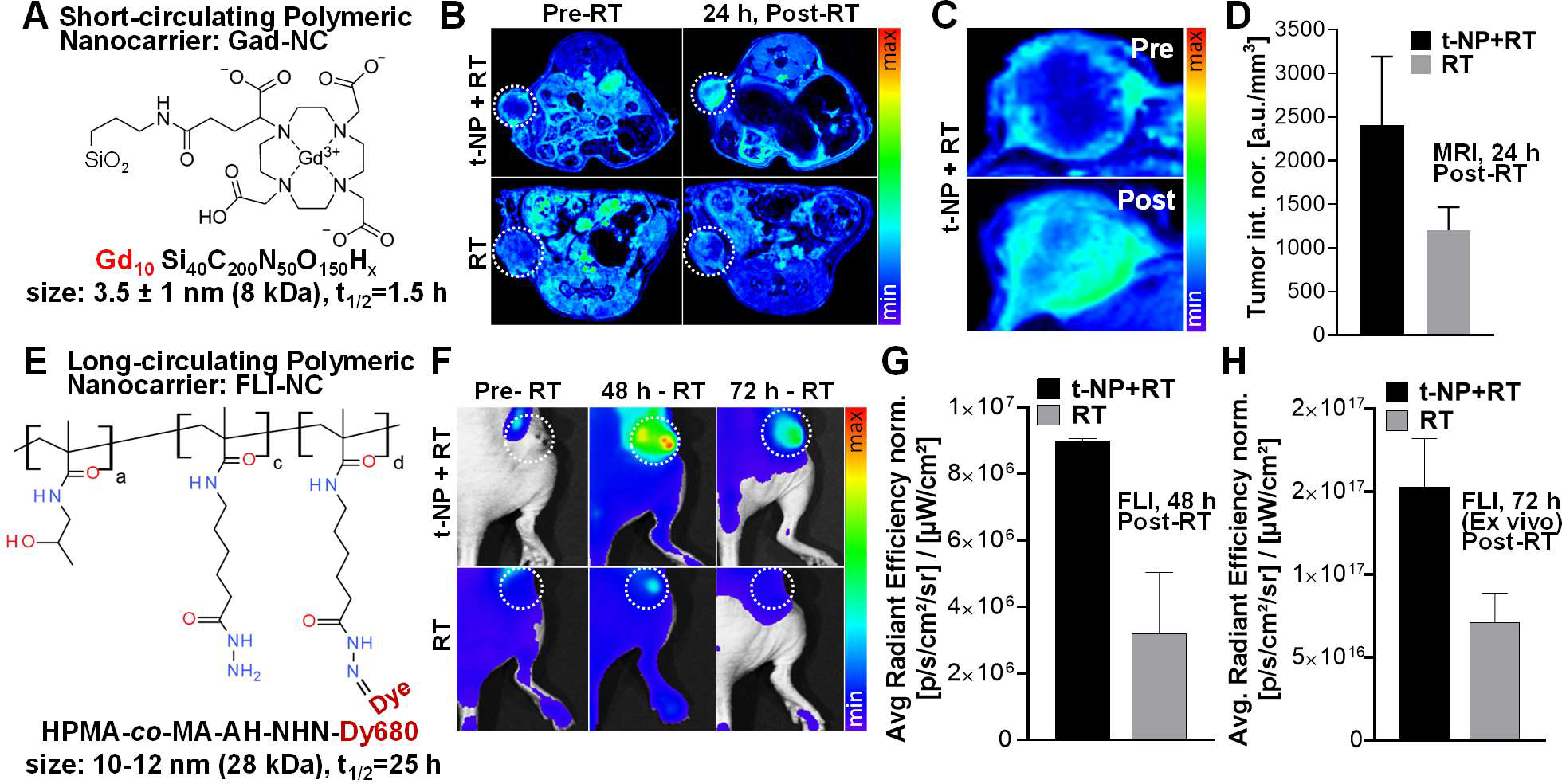
Enhanced tumor-specific drug delivery in the *h*-PDAC model. To trigger tumor-specific drug delivery post-vascular disruption using t-NP and rT, two nanocarrier formulations with short and long circulating properties were employed. **A**. Gadolinium-based nanoformulations (Gad-NC) were used to perform tumor uptake studies pre- and post-tumor vascular disruption. **B**. T_1_-weighted MRI demonstrated high uptake in the t-NP+RT treated group compared to the RT only group. **C-D**. Magnified image shows the intratumoral distribution of the nanocarrier towards the core of the tumor post-RT treatment. Upon further quantification, a twofold difference between the t-NP treated *vs.* non-treated cohorts was evident. **E-F**. A long-circulating polymeric nanocarrier of HPMA coupled to a fluorescent dye (FL-NC) was administered to mice bearing a Capan-1 pancreatic tumor, and fluorescence imaging were performed. A longitudinal accumulation of FL-NC in higher amounts was observed in the t-NP+RT-treated tumors, both at 48 h and 72 h (ex *vivo).* **G-H**. Further quantification of fluorescent signals demonstrated a >two-fold increase in the accumulation of FL-NC in *h*-PDAC tumors. All values were normalized to its respective standards, and four different control conditions were used.

In *h*-PDAC, T_1_-weighted MRI demonstrated an overall increase in Gad-NC accumulation in the ‘tumor vascular modulated’ t-NP+RT group at 24 h post-RT (**Fig. 5B**). Intratumoral uptake examined across the reconstructed 3D slices of the tumor MRI show prominent distributions of Gad-NC in the periphery as well as the core of the tumor, as opposed to the RT-only group which demonstrates contrast enhancement mostly in the tumor periphery regions (**Fig. 5C**). MRI-based quantification confirmed a twofold difference in the accumulation of Gad-NC in the t-NP+RT *vs*. RT-only groups (**Fig. 5D**). To further assess the dynamics of tumor accumulation for prolonged periods of time post-RT, an HPMA-based fluorescent nanocarrier (FL-NC) was used as a model nanocarrier (**Fig. 5E**). HPMA-based drug delivery systems are ideal for EPR-mediated drug targeting due to their prolonged circulation half-life (t_1/2_=25 h), biocompatibility, and non-immunogenicity. ^38, 48^ Both qualitative and quantitative results demonstrated more than two-fold increase in FL-NC accumulation in the tumors that underwent vascular modulation during t-NP+RT treatment, compared to the RT-only treatment (**Fig. 5F-G**). *Ex vivo* tumor samples at 72 h post-RT confirmed the differences in the tumor accumulation of FL-NC in an *h*-PDAC tumor model (**Fig. 5H**).

### Assessment of tumor blood vessel functionality post-modulation

Tumor vascular modulation results in apparent changes in the permeability (*K*_*trans*_), extravascular back-flux (*K*_*ep*_), and extravascular extracellular volume fraction (*V*_*e*_) (**Fig. 6A**). DCE-MRI studies using Gad-NC at 24 h post-RT demonstrated an increase in the tumor vascular permeability (*K*_*trans*_) for nanoparticle combined radiation group compared to the radiation-only group (**Fig. 6B**). Furthermore, the *K*_*ep*_ values indicate a low-extravascular back-flux into the blood plasma, further confirming the tumor/interstitial retention of Gad-NC. Similarly, the decrease in the extracellular extravascular volume fraction (*V*_*e*_) indicates perfused Gad-NC that has been taken up by the cancer cells - a scenario that ideally depicts the tumor uptake of chemo or nano therapeutics. Quantitative measurements confirmed apparent differences in the *K*_*trans*_, *K*_*ep*_ and *V*_*e*_ parameters among the t-NP+RT *vs.* RT-only tumors (**Fig. 6C-E**). These changes were consistent with the increase in tumor-specific drug delivery in *h*-PDAC (**Movie S1**, **S2**). Moreover, the acute changes in tumor permeability were measured across the different tumor slices, i.e., from the periphery to the core (**Fig. 6F**). There is an overall increase in tumor vascular permeability from core to periphery, however, in almost all the tumor slices, t-NP+RT demonstrate high Gad-NC accumulation. The qualitative evidence further confirmed this improved Gad-NC distribution (**Fig. 6G**). In short, both the MR and FLI studies explicitly confirmed the improved tumor uptake of nanocarriers or model drug delivery systems and substantiated the concomitant changes in the tumor vascular parameters such as *K*_*trans*_, *K*_*ep*_, and *V*_*e*_ that led to improved and targeted drug delivery in *h*-PDAC.

**Figure 6:**
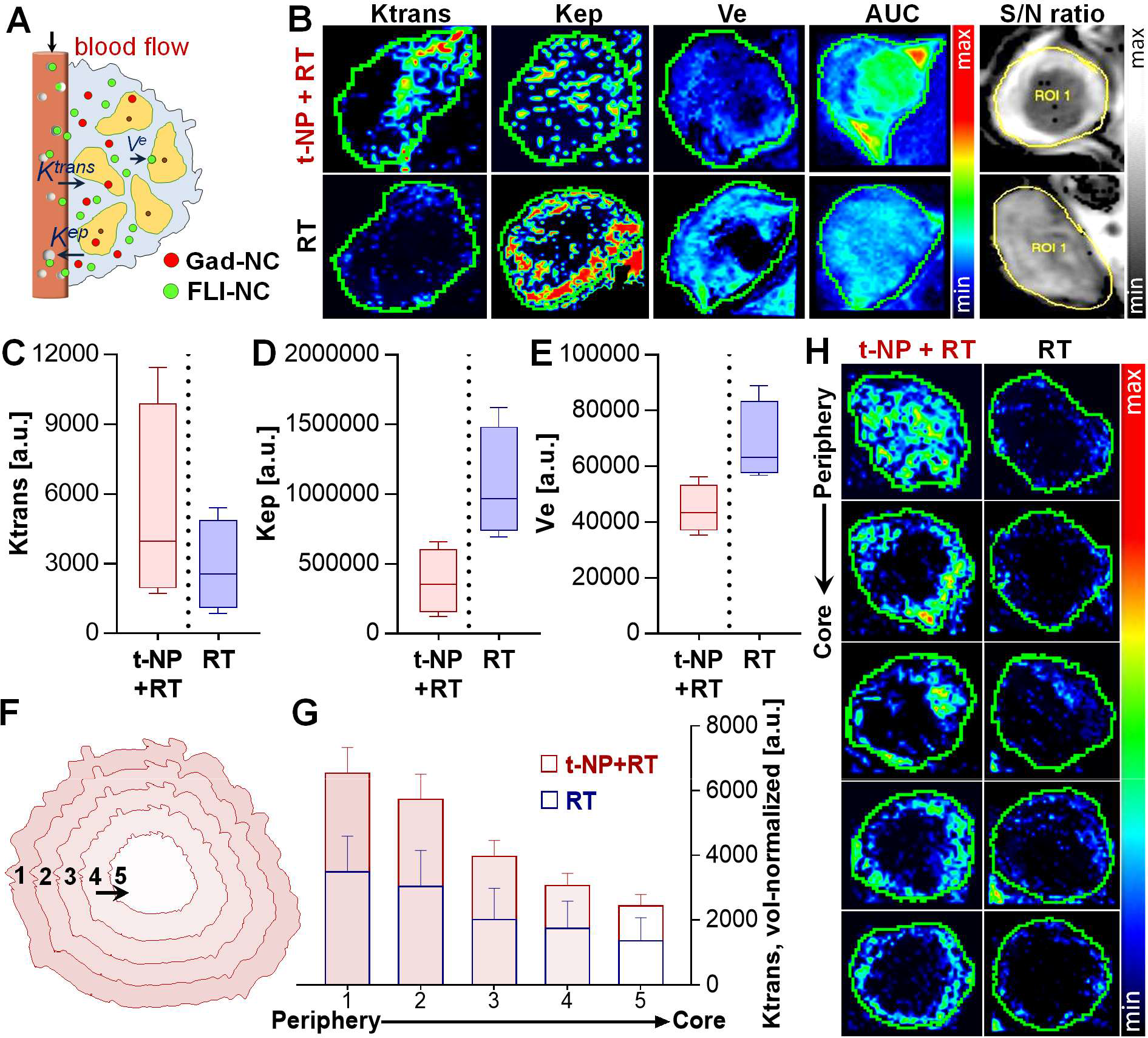
Assessing dynamic changes post tumor vascular modulation. **A-B**. Dynamic changes in the tumor vascular parameters *K_*trans*_* (transendothelial permeability), *K*_*ep*_ (extravascular back-flux), and Ve (extravascular extracellular volume fraction) were measured using DCE-MRI after *i.v*.-administration of Gad-NC to *h*-PDAC tumor-bearing mice. The measurements were carried out and compared to the t-NP+RT and RT group, along with other non-treated control groups. DCE-MRI studies displayed a qualitative increase in the tumor vascular permeability (*K*_*trans*_) following tumor vascular modulation, and associated decrease in *K*_*ep*_ (back-flux into the plasma) and *V*_*e*_ (the extravascular extracellular volume fraction) parameters. **C-E**. Further changes induced by t-NP+RT treatment was quantitatively measured by an increase in *K*_*trans*_ (permeability); and a concomitant decrease in *K*_*ep*_ (retention) and *V*_*e*_ (uptake) - prototypical responses of an anti-vascular treatment. Discontinuous lines relate to the ‘tumor vascular modulation’ by t-NP+RT treatment *vs.* ‘no modulation’ in RT-only treatment. **F-G**. Intratumoral changes in vascular permeability from the tumor periphery to the core (in 3D) was manually segmented, thus covering the entire tumor. The corresponding *K*_*trans*_ plots show definite improvement in the endothelial permeability from the tumor periphery to the core in vessel-modulated t-NP+RT cohorts, compared to the RT-only cohorts. **H**. Two-dimensional tumor slices were analyzed, and *K*_*trans*_ changes were further confirmed qualitatively.

## CONCLUSIONS

EPR forms the basis for tumor accumulation in the case of most passively-targeted clinical nanomedicine formulations. ^37, 52–53^ Accumulation of nanodrugs in the tumor is related to increased hyperpermeability of the tumor neovasculature. ^38, 42, 52, 54^ For instance, a clinical study tested PEGylated liposomes of doxorubicin in squamous lung carcinoma, head and neck cancer, and breast cancer - tumor types with varying degrees of vascular permeability. ^55^ Interestingly, the highest tumor accumulation of (doxorubicin-containing) liposomes was found in the head and neck cancer (33 ± 16%ID/kg), and intermediate accumulation in the lung adenocarcinoma tumors (18 ± 6% ID/kg), and relatively low accumulation in breast cancer patients (5 ± 3% ID/kg). ^55^ These results suggest that high-EPR (or increased tumor permeability) may translate to improved nanodrug uptake and consequent therapeutic benefit.

Tumor permeability is variable across various tumor types, and even within a single tumor type, the heterogeneous distribution of permeability is ubiquitous. ^17, 56^ In slow-growing preclinical tumor models (that which resembles human-like tumors), intact and mature tumor blood vessels with high neoangiogenic receptor expression are commonly observed. ^38^ For instance, PDAC tumors have less leaky neovessels and display comparably low-EPR; however, it has profound expression integrins (or RGD) on its tumor endothelium. This opportunity was explored for targeting RGD in *h*-PDAC tumor endothelium. Experimental findings from our study show the first-proof-of principle demonstrating radiation-induced tumor vascular alteration that can enhance tumor-specific drug delivery in a human pancreatic tumor model. The proposed strategy may also be applied to other non-resectable or intractable tumors that respond poorly to standard clinical therapies.

### Statistical Analysis

Data are expressed as a mean ± standard deviation or standard error unless otherwise indicated. Statistical analyses and graphs were carried out using Prism (GraphPad Software, Inc., La Jolla, CA, USA). The unpaired, two-tailed Student’s t-test was used to determine significance between an experimental group and a control group with a value of P≤0.05 considered significant. For comparisons between multiple groups, simple one-way ANOVA test was used. Kaplan-Meier plots with log-rank (Mantel-Cox) tests were used for survival studies.

## ACKNOWLEDGMENTS

We greatly acknowledge the efforts by histology core at Brigham and Women’s Hospital and Harvard Medical School, and the TEM core facility at Harvard Medical School. We thank Anirudha Karve (DFCI) for his technical assistance with small animal radiation therapy and F Pelascini (CRITT Matériaux Alsace) for technical assistance with epoxy-embedding and LIBS imaging of biological tissue samples.

## ASSOCIATED CONTENT

Supplementary Information

